# Low atmospheric CO_2_ levels induce nocturnal carbon accumulation in the lycophyte genus *Isoëtes*

**DOI:** 10.1101/820514

**Authors:** Jacob S. Suissa, Walton A. Green

**Affiliations:** Department of Organismic and Evolutionary Biology, Harvard University, Cambridge, MA; Arnold Arboretum of Harvard University, Boston, MA

**Keywords:** CAM evolution, Isoetaceae, CO_2_ manipulation, atmospheric CO_2_, *Isoëtes engelmannii*, *Isoetes tuckermanii*, quillwort, isoetid physiology, aquatic CAM

## Abstract

**Background and Aims:** Crassulacean Acid Metabolism (CAM) is an adaptation to increase water use efficiency in dry environments. Oddly, similar biochemical patterns occur in the submerged aquatic lycophyte genus *Isoëtes*. It has long been assumed that CAM-like nocturnal carbon accumulation in aquatic plants is an adaptation to low daytime carbon levels in aquatic ecosystems, but this has never been directly tested.

**Methods:** To test this hypothesis, populations of terrestrial *Isoëtes engelmannii* and *I. tuckermanii* were grown in climate-controlled chambers and starved of atmospheric CO_2_ during the day while pH was measured for 24-hours.

**Key results:** We demonstrate that terrestrial plants exposed to low atmospheric CO_2_ display diel acidity cycles similar to those in both xerophytic CAM plants and submerged *Isoëtes*.

**Conclusions:** CO_2_ starvation induces CAM-like nocturnal carbon accumulation in terrestrial *Isoëtes*, substantiating the hypothesis that carbon starvation is a selective pressure for nocturnal carbon accumulation in *Isoëtes*. Furthermore, while aquatic carbon levels undoubtedly promote nocturnal carbon accumulation in extant *Isoëtes*, the induction of this behavior in *terrestrial* plants suggests a possible earlier terrestrial evolution of this metabolism in Isoetalean ancestors in response to low *atmospheric* CO_2_ levels. We both provide support for a long-standing assumption about nocturnal carbon accumulation *Isoëtes* as well as suggest an earlier evolution of this behavior.

## INTRODUCTION

Metabolic shifts play an essential role in the survival of plants in extreme habitats. Many desert plant species, for example, minimize water loss by temporally segregating the light and dark reactions of photosynthesis. This temporal segregation allows plants to limit water loss during the day, while incorporating CO_2_ into 4-carbon acids at night for subsequent daytime fixation (Kluge and Ting 1978; Sipes and Ting 1985; Lüttge 2004). This well-known behavior is known as Crassulacean Acid Metabolism (CAM) and leads to noticeable diel cycles in pH and acidity of photosynthetic organs. This metabolism is critical for the success of many xerophytic plants and is regarded as one of the most important photosynthetic adaptations to dry environments (Kluge and Ting 1978). Although CAM in xerophytic plants allows for high water use efficiency in dry environments, the pathway behind this metabolism (C4, Hatch/Slack/Korshak pathway) is fundamentally a carbon concentrating mechanism which increases the selectivity of rubisco by altering the CO_2_:O_2_ ratio within the chloroplast (Keeley and Rundel 2003; Edwards 2019).

In 1981, diel acidity cycles like those observed in CAM plants were first observed in the aquatic lycophyte genus *Isoëtes* (Keeley 1981). This metabolism was called ‘aquatic CAM’ to highlight its similarity to (xerophytic) CAM acidity cycles. Here, we use the term ‘nocturnal carbon accumulation’ (NCA) to emphasize the differences between the behavior described in *Isoëtes* and that in xerophytic CAM plants. Further investigations of NCA in *Isoëtes* have demonstrated a high degree of plasticity. For example, *Isoëtes howellii* exposed to atmospheric CO_2_ levels while growing terrestrially tends not to accumulate carbon nocturnally, while *I. karstenii* accumulates carbon regardless of its environment, demonstrating that this behavior can be facultative or constitutive (Keeley 1998; Keeley and Rundel 2003). Since CAM is first and foremost a carbon concentrating mechanism, it has long been assumed that NCA in *Isoëtes* is a response to low daytime carbon availability in aquatic ecosystems–but this assumption has hitherto remained largely hypothetical and correlative (Keeley 1982, 1983, 1998; Keeley and Bowes 1982; Keeley and Rundel 2003). Moreover, it is suggested that the selective pressure for the evolution of NCA in *Isoëtes* is hypocarbia in eutrophic and oligotrophic lakes (Keeley and Rundel 2003)–although an alternative possibility is that NCA is an adaptation that evolved early in this lineage in response to low *atmospheric* CO_2_ levels in terrestrial ancestors of *Isoëtes*, and was inherited and co-opted by *Isoëtes* in aquatic environments where daytime CO_2_ is limiting and diffusion rates for CO_2_ are four orders of magnitude slower than in air.

To test the hypothesis that NCA is a direct response to carbon starvation, we grew terrestrial plants of two *Isoëtes* species in environmentally controlled growth chambers, starved plants of atmospheric CO_2_ during the day and sampled leaf pH from multiple plants for 24-hours. We demonstrate that, diurnal atmospheric hypocarbia induced diel acidity cycles in terrestrial *Isoëtes engelmannii*, like those observed in xerophytic CAM plants and submerged aquatic *Isoëtes* species (Fig. 4c). This provides direct evidence to support the hypothesis that CO_2_ limitation promotes CAM-like NCA in extant *Isoëtes* species–as previously hypothesized (Keeley and Bowes 1982; Keeley 1983; Keeley and Rundel 2003). Moreover, the induction of NCA in terrestrial *Isoëtes* by atmospheric CO_2_ starvation may support an alternative hypothesis that CAM-like photosynthesis evolved early in terrestrial Isoetalean relatives in the Carboniferous in response to globally low atmospheric CO_2_ levels, as opposed to a more recent aquatic evolution (Keeley 1998).

**Figure 1:**
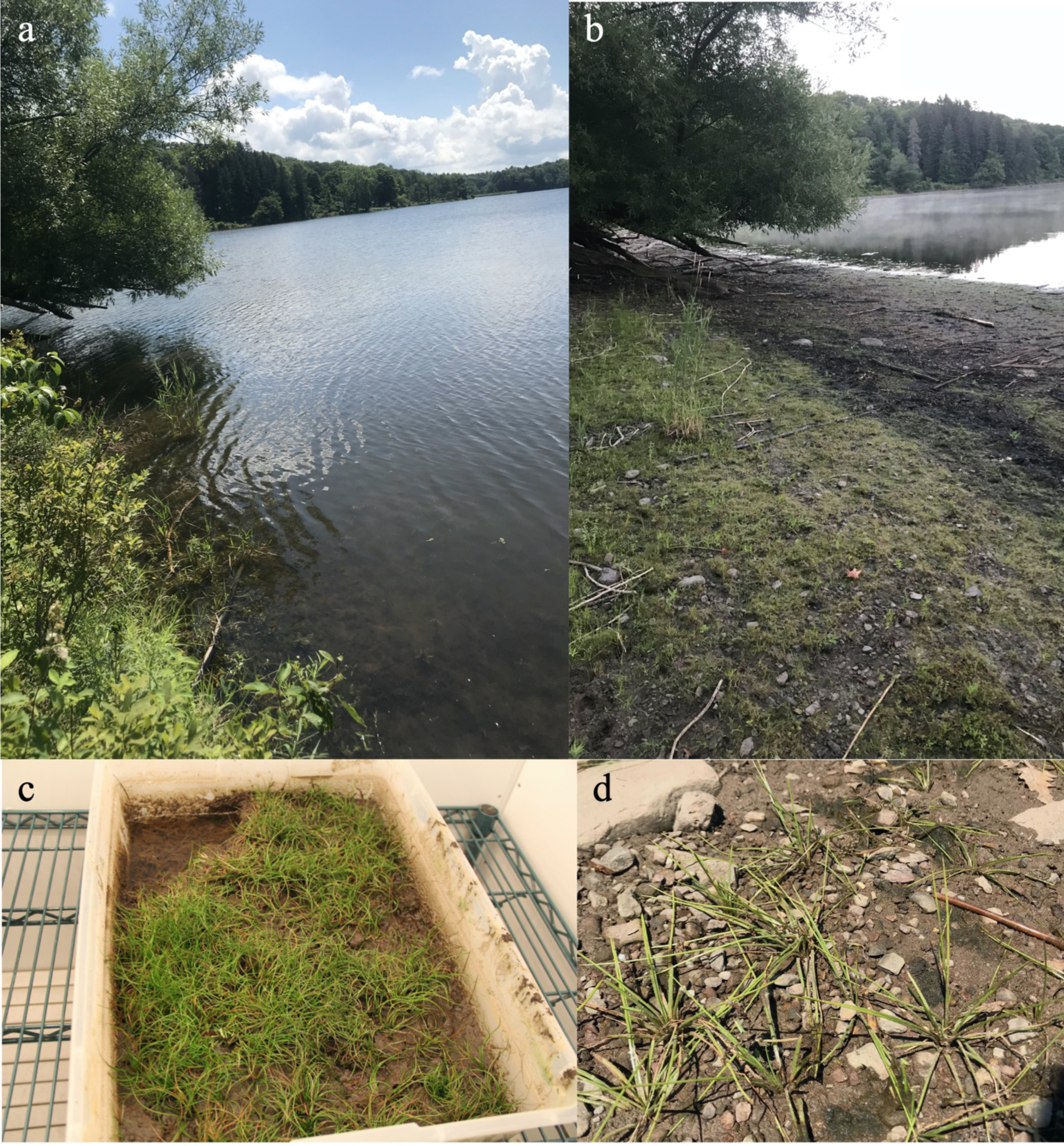
a: Lake Myosotis study site during average water levels for most of the year. b: Lake Myosotis in July when water levels drop. c exposed plants of I. tuckermanii in lab conditions. d: exposed plants of I. engelmanii in situ.

**Figure 2:**
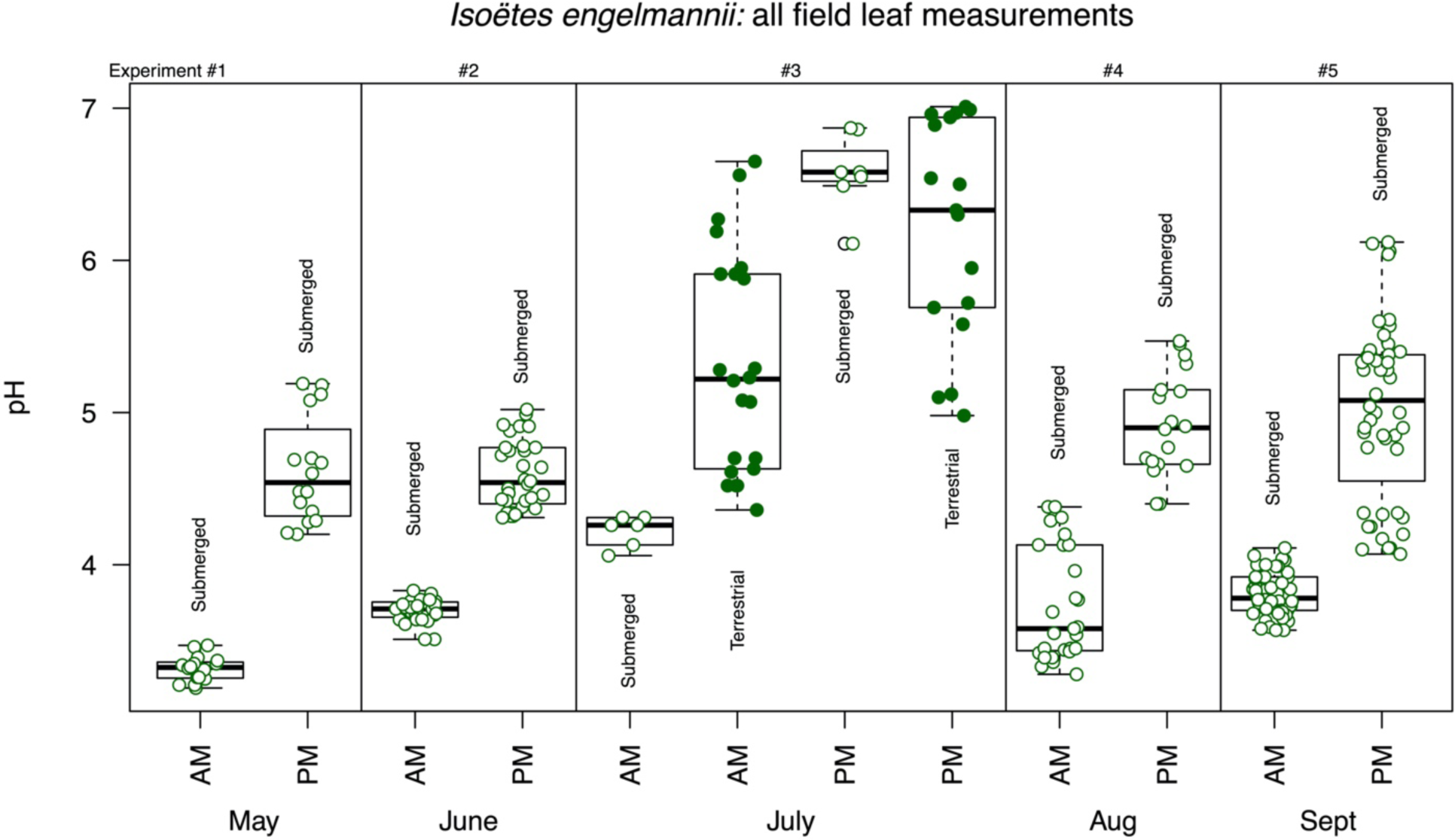
Field experiments. Diel pH measurements of leaves from field grown plants throughout the summer months. In July some plants were growing terrestrially and did not show significant differences in pH. Each point represents a single pH measurement. A total of 0.2-0.5g of leaf material was pooled from 5-6 individuals and separated into 3-8 distinct samples. During September more than two time points were measured throughout the day (see Fig. 3).

**Figre 3:**
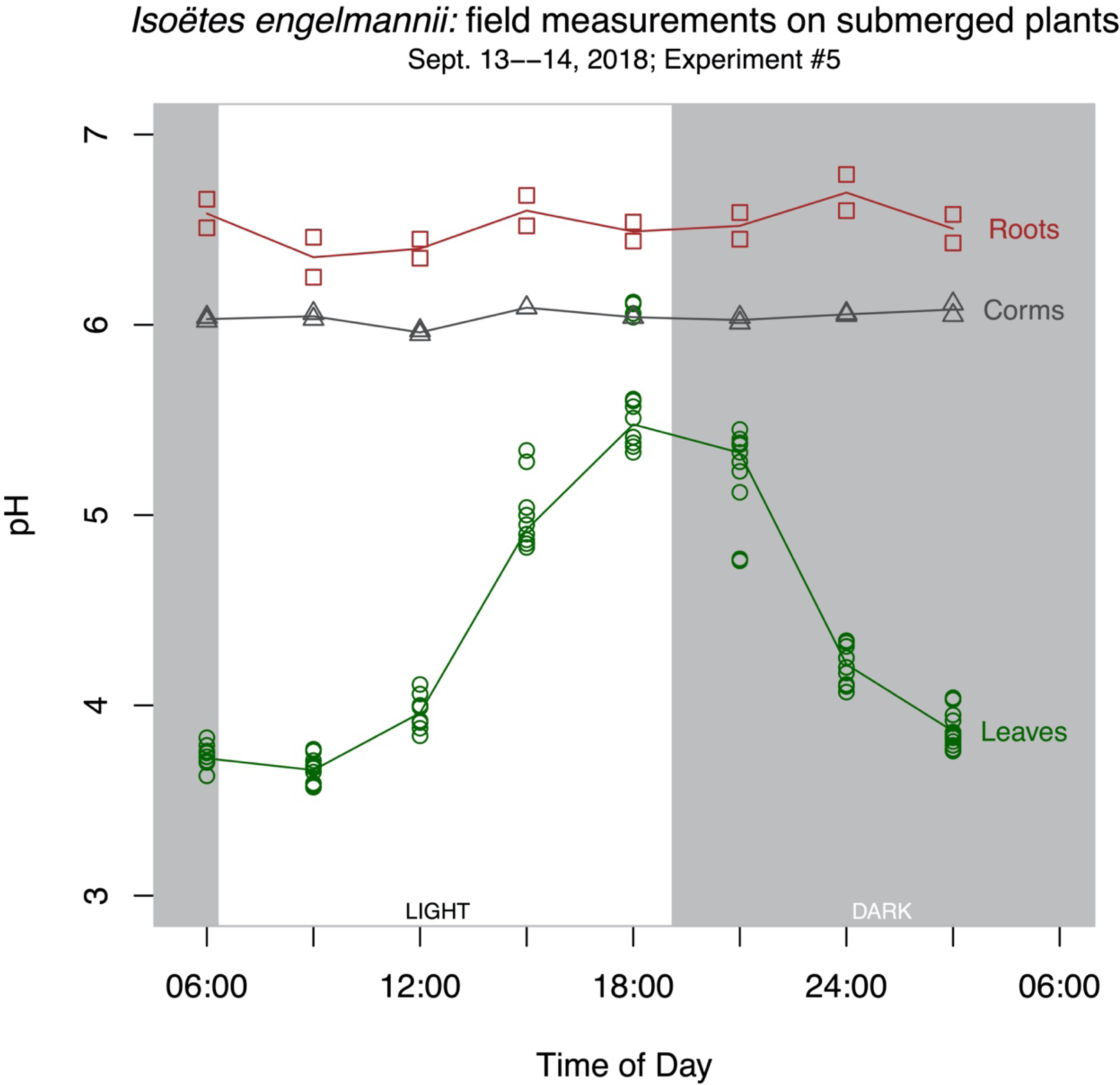
Field experiments. pH measurements of leaves, corms and roots, for 24-hours in September. A clear cycle in pH is demonstrated in leaves while corms and roots show no diel change. Each point represents a single pH measurement of 3-8 samples comprised of 0.2-0.5g of pooled leaves from 5-6 individuals, pH was measured twice on each sample.

**Figure 4:**
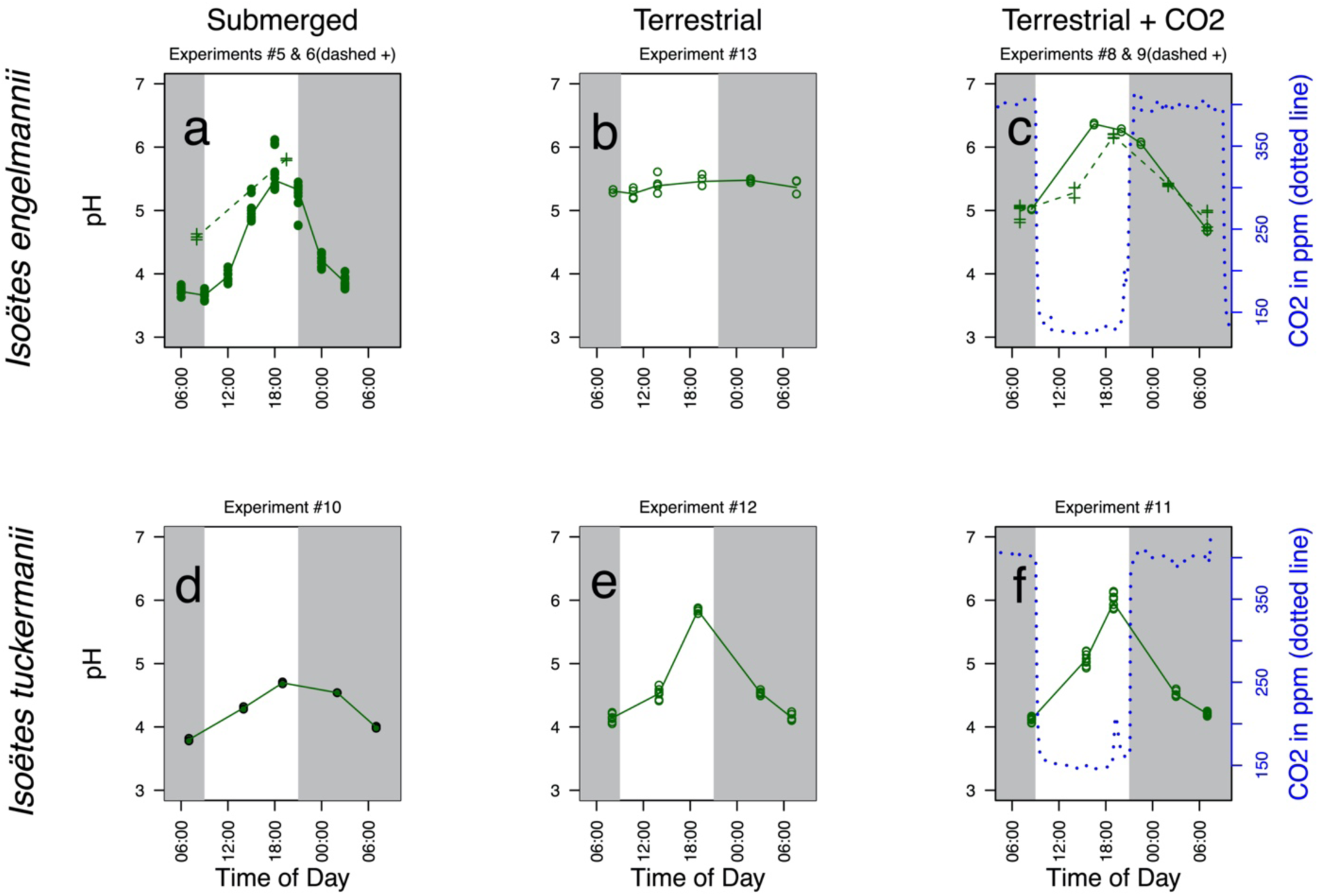
Laboratory experiments. Leaf pH measurements of I. engelmanii and I. tuckermanii under, a, d. submerged ambient CO2 levels; b, e. terrestrial ambient atmospheric CO2 levels; and c, f. terrestrial CO2 manipulation. A clear diel change in pH was not observed in I. engelmanii when emergent but was induced upon CO2 starvation. Solid circles represent submerged plants and open circles represent terrestrial plants. Each point represents a single pH measurement of 2-3 samples comprised of 0.2-0.5g of pooled leaves from 2-4 individuals, pH was measured twice on each sample. *Circles in pane a are measurements from the field, crosses represent lab experiments. In pane c two separate 24-hour experiments were conducted (crosses and circles).

## MATERIALS AND METHODS

### Field experiments

Field Experiments: The Edmund Niles Huyck Preserve is located in Rensselaerville, NY, at 425–525m elevation. Plants of *Isoëtes engelmanii* were studied from May–September in a population on the north shore of Lake Myosotis (Russell 1958), 42°31’26.8” N 74°9’7.2” W. Three to eight samples each composed of leaves, corms, or roots, from 5–6 plants were collected five times during the summer of 2018 from May–September (Fig. 2). During May, June, and July, samples were collected only in the morning at 0600 and in the evening at 1800; in August, samples were collected every three hours from 0600–1800, and in September collections were made every three hours for a full 24 hours (1800–1800). During all experiments plants were cleaned of algae and soil in DI water in the lab, which was located 1km from the study site.

In the lab, leaves, corms, and roots were separated using scalpels and forceps and organs were washed and blotted dry. At each time point, leaves from 5–6 individuals of *I. engelmannii* were randomized and separated into 3–8 distinct samples, depending on the amount of material available. Each of the individual samples was composed of 0.2–0.5g of leaf, corm, or root tissue. Tissue from each of the 3–8 samples was macerated using a plastic dowel in a 1.7mL Eppendorf tube, and 0.5mL of DI H2O was added to each sample. Samples were then resuspended by mixing using a vortex mixer at maximum intensity for 10 seconds, followed by centrifugation at 10,000 rpm for 10 seconds. The supernatant was carefully extracted using a pipette, and placed on the Horiba LAQUAtwin pH-22 meter, for pH readings. The pH was measured twice from each sample to ensure accuracy of the reading. The pH meter was cleaned with deionized water and dried between each sample. The pH meter was recalibrated between each time point using pH 4 and 7 standardized buffers. These measurements were made instead of the historically used acid titration method because of the ease of use of handheld pH meters and the small amount material needed to produce accurate measurements. The main differences between measuring acid levels using pH meters compared to acid titration is that pH measured from two solutions with the same amount of acid can be different if they have different buffering capacities (Sadler and Murphy 2010). Even though there are difficulties with comparing pH and titrant concentration with weak acids in solutions of different concentrations, one can still relate the two by using the Henderson-Hasselbalch equation (Sadler and Murphy 2010). We make the assumption that the buffering capacity of leaves of *Isoëtes* does not substantially change over the course of a 24-hour period, which justifies the use of a pH meter to quantify change during the diel cycle. If leaves from different species had been compared, buffering capacity could potentially be different and pH measurements potentially harder to compare between samples without quantitative titration.

### Laboratory experiments

*Isoëtes engelmannii* specimens collected in the field throughout summer 2018 in Lake Myosotis were cultivated in growth chambers in the Weld Hill Research Building of the Arnold Arboretum of Harvard University. In addition, populations of *I. tuckermanii* already growing in the greenhouse (collected from the NW shore of Lake Mattawa, in Orange, MA, 42°34’11.9” N 72°19’34.1” W), were used in the growth chamber experiments. For 2 months, plants were grown fully submerged at ambient CO_2_ levels (∼400ppm), at 20°C, with a 12-hour photoperiod, 15µmolm^-2^ photosynthetically active radiation. Submerged pH was measured on a diel cycle following the field protocol (see above). Containers were then drained of water, and the plants were allowed to acclimate for 1–3 days. While plants were terrestrial under ambient CO_2_ levels, pH was measured every 3–5 hours for 24 hours following the aforementioned protocol. Plants were then moved to a growth chamber set to a diel cycle of atmospheric CO_2_: 100ppm of CO_2_ during the 12-hour photoperiod (the minimum obtainable in the chamber) and 400ppm (equivalent to ambient atmospheric levels) during the dark period. Temperature and light intensity were not changed from the ambient CO_2_ conditions. Starting at 0700, plants were harvested every 3–5 hours for 24 hours. At each time point leaves from 2–4 individuals were randomized and separated into 2–3 distinct samples comprised of 0.2–0.5g of leaf tissue and pH was measured following the field protocol.

### Statistics

To quantify the similarity in the diel cycle of acidity between two experiments, we employed two statistical methodologies. First, we applied the Brown-Forsythe test for the equality of variance (Brown and Forsythe 1974). Second, we calculated the overlap in tail probabilities of the probability distribution of the Maximum Value Unbiased Estimator (MVUE) for the maximum diel range. Neither of these approaches is optimal, but each provides some indication of the likelihood that the observed diel cycles are significantly different at the indicated probability level. In the case of the Brown-Forsythe test, this is a conventional p-value; in the alternate MVUE approach, it is a (one-tailed) probability that the larger diel range would be observed, given the distribution observed in the sample with a smaller diel range. All manipulations and the pairwise comparisons between all experiments are provided and fully documented in the code supplement. In the text, the results of a statistical comparison between two experiments is shown in the format x:y where the comparison is between experiment x and y, prefixed by the type of test (Brown-Forsythe or MVUE). So ‘Brown-Forsythe (5:6), p=0.03’ means that the Brown-Forsythe test rejected the null hypothesis that experiments 5 and 6 had equal variances at the 5% level (but not at the 1% level). Experiment numbers referred to in the text correspond with those shown in the figures.

## RESULTS

### Field experiments

In our first series of field measurements, from May to September, we measured morning and evening pH in *Isoëtes engelmannii* leaves, roots, and corms, monthly. (Fig. 1a; Fig. 2). We found that in the field, submerged plants of *I. engelmannii* accumulated carbon on a diel cycle (Fig. 2; Fig 3). The mean morning pH of the leaves throughout the summer was 3.71 and the mean evening pH was 4.90 (Fig. 2). The pH of non-photosynthetic organs (roots and corms) did not change on a diel cycle, showing that the NCA was restricted to the leaves (Fig. 3); corms: mean morning pH 5.94, mean evening pH 5.97; roots: mean morning pH 6.32, mean evening pH 6.36 (Fig. 3). Upon recession of the shoreline in July, many of the plants were left growing terrestrially, exposed to atmospheric conditions (Fig. 1). In the plants growing terrestrially, pH had higher variability in both the morning and the evening (Fig. 1d; Fig 3), and the variance of terrestrial measurements was significantly higher than submerged values throughout the summer (p<0.001 based on a Brown-Forsythe test for equality of variance and tail probabilities of a minimum variance unbiased estimator of maximum diel range (MVUE) (3:1–5)). Prior observations of *I. engelmannii* additionally also suggest that this species accumulates carbon nocturnally only when submerged (Keeley 1998).

### Lab experiments

For laboratory manipulations, we collected *I. engelmannii* from the same population measured in the field as wells as specimens of *I. tuckermanii*, a related species that generally grows completely submerged. All plants collected were brought into the lab and cultivated in climate-controlled growth chambers. Under ambient CO_2_ levels the diel pH variation of submerged individuals of *I. engelmannii* was similar to submerged individuals in the field (Fig. 4a) (MVUE (5:6) p=0.01; Brown-Forsythe (5:6), p=0.03); mean leaf morning pH was 4.58, and mean evening pH was 5.81. Submerged plants of *I. tuckermanii* had a qualitatively less dramatic diel shift in pH, compared to *I. engelmannii* (MVUE (5:10) p<0.001, (6:10) p=0.04; Brown-Forsythe (5:10) p=0.28, (6:10) p=0.01), but still experienced a diel fluctuation (Fig. 4a, 4d); mean morning pH was 3.89, and mean evening pH was 4.70. When emergent, as expected from the field experiments, individuals of *I. engelmannii* no longer displayed a diel shift in pH (Fig 4b; Fig. 2) (MVUE (5:13) p<0.001, (6;13) p<0.001; Brown-Forsythe (5:13) p<0.001, (6:13) p=0.53); mean morning pH was 5.31, mean evening pH was 5.46. Experiment 6 was only comprised of 2 time points, thus statistical support for determining differences is lower when compared to other experiments (**Supplemental table**). *Isoëtes tuckermanii*, however, showed a more dramatic diel change in pH, when emergent compared to when it was submerged (MVUE (10:12) p=0.001; Brown-Forsythe (10:12) p=0.02) (Fig. 4e); mean morning pH was 4.15, mean evening pH was 5.84. These results demonstrate different NCA behavior in these two species, as has been previously documented in *Isoëtes* (Keeley 1998). *I. engelmannii* induces nocturnal carbon accumulation when submerged but not when emergent, while *I. tuckermanii* demonstrated constitutive NCA, with similar diel variation in pH irrespective of water depth, and only a slight increase in NCA when emergent compared to when submerged.

We next grew the plants with diurnal CO_2_ starvation and nocturnal enrichment to mimic the CO_2_ conditions in a eutrophic lake. *Isoëtes engelmannii* individuals grown under these conditions again showed a diel cycle in pH similar to that observed in the submerged specimens in lab and field, but different from emergent plants in the field and lab (MVUE (5:8) p=0.01, (6:8): p=0.17, (5:9) p<0.001, (6:9) p=0.34); Brown-Forsythe (5:8) p<0.001, (6:8) p=0.29, (5:9) p<0.001, (6:9) p=0.72) (Fig. 3a; 3c). In two independent experiments with multiple replicates per time point, we measured a mean morning pH of 4.69, mean evening pH was 6.37 (Exp. 8); mean morning pH 4.85, mean evening pH 6.18. The magnitude of pH change in the terrestrial plants grown under diurnal CO_2_ starvation was similar to that in submerged plants in the field experiments, though slightly dampened (Fig. 3; Fig 4c). This may be due to the fact that predawn CO_2_ concentrations in the field in eutrophic lakes and ponds can exceed 2500ppm (Keeley and Bowes 1982), and the maximum CO_2_ enrichment during our experiment approximated ambient atmospheric concentrations of 400 ppm (Fig. 3c).

When grown under these manipulated CO_2_ conditions, *I. tuckermanii* continued to show nocturnal carbon accumulation (Fig. 1c; Fig. 3f) with a mean morning pH of 4.12, and a mean evening pH of 5.98. The pH fluctuation of *I. tuckermanii* during diurnal hypocarbia did not differ from that of the terrestrially growing plants in ambient CO_2_ (MVUE (11:12) p=0.02; Brown-Forsythe (11:12) p=0.38) (Fig. 3e), suggesting that this species is obligate in its NCA. Unlike *I. engelmannii*, the individuals of *I. tuckermanii* we examined had no stomata, a possible explanation of this constitutive behavior. These results provide a possible explanation for the plasticity of NCA in the genus *Isoëtes* observed in the past (Keeley 1982, 1998). Moreover, we show that laboratory-based experimental modification of natural conditions can replicate field conditions and that carbon starvation imposed on terrestrial plants in the lab produces a diel change in pH comparable to the change observed in submerged aquatic plants.

## DISCUSSIONS

In this study, we demonstrate that nocturnal carbon accumulation–like that in xerophytic CAM plants and submerged *Isoëtes*–can be induced in terrestrial *Isoëtes* by limiting day-time atmospheric CO_2_ levels. While it has been hypothesized for over 40 years, these findings provide new and substantial evidence to support the hypothesis that carbon limitation induces nocturnal carbon accumulation in *Isoëtes*. Furthermore, because low *atmospheric* CO_2_ levels can induce NCA in terrestrial *Isoëtes* in the same way as low *aquatic* CO_2_ levels do, it seems possible that low atmospheric CO_2_ levels may have been important in the early evolution of this metabolic pathway in the Isoetalean lineage.

Prior discussions of the adaptive significance of NCA in *Isoëtes* have focused on aquatic CO_2_ limitation as a selective force (Keeley 1981, 1982, 1983, 1996, 1998; Keeley and Bowes 1982; Keeley *et al*. 1983; Keeley and Rundel 2003). Our experiments now show that terrestrial carbon limitation has the same effect. Low atmospheric CO_2_ was a notable feature of the Carboniferous (Beerling 2002; Van Der Meer *et al*. 2014), a time when Isoetalean lycopsids formed a substantial part of the terrestrial flora. In the Carboniferous atmosphere, a carbon concentrating mechanism that increased carbon gain and minimized the photorespiratory loss of energy in a high oxygen/low CO_2_ atmosphere would have been particularly advantageous. Given that we induced NCA in terrestrial *Isoëtes* (Fig 3c), and that this behavior is widespread across the genus (Keeley 1982, 1998), it may be possible that NCA evolved in the early Isoetalean lycopsids during the low atmospheric CO_2_ conditions of the Carboniferous period. If so, NCA in extant aquatic *Isoëtes* would represent an exaptation of a much older metabolic behavior rather than a more recent adaptation to low aquatic CO_2_ levels. An alternative interpretation of our results is that low atmospheric CO_2_ levels mimic the extant aquatic environment, meaning that NCA may still be an adaptation to the recently inhabited aquatic ecosystem. While this remains a possibility, what is known of the ecology and evolution of the Isoetalean lineage suggests that atmospheric rather than aquatic limitation of CO_2_ may have contributed to the evolution of NCA in Isoetales. Whether nocturnal carbon accumulation in *Isoëtes* evolved earlier or later–in response to atmospheric or aquatic CO_2_ levels–more consideration should now be given to the selection pressures on net assimilation and the source of carbon limitation leading to the evolution of NCA.

Our results provide direct evidence for the induction of NCA in extant plants by carbon limitation, exact determination of the evolutionary timing of NCA in *Isoëtes* requires more complete consideration of the temporal variation and selective pressures of local aquatic and global atmospheric CO_2_ levels, as well as the phylogenetic distribution of this behavior in extant *Isoëtes* species. Examining a broader range of *Isoëtes* species and using phylogenetic comparative methods is likely to show multiple independent appearances of constitutive and inducible NCA throughout the extant genus (Keeley 1998; Keeley and Rundel 2003), as well as in other plants conforming to the Isoetid habit, just as CAM has been found to evolve many times in vascular xerophytes (Edwards and Ogburn 2012; Hancock *et al*. 2019).

## Conclusions

Knowledge of carbon uptake and photosynthetic strategies in an evolutionarily distinct lineage such as the Isoetaceae represents an important step forward in understanding the evolution of carbon concentrating strategies. This study is the first to provide direct evidence that CO_2_ starvation induces CAM-like nocturnal acid accumulation in the aquatic macrophyte *Isoëtes*. Further, the induction of NCA in *Isoëtes* by low atmospheric CO_2_ levels may suggest that NCA evolved in early in the Isoetaceae as an adaptation to low atmospheric CO_2_ levels in the Carboniferous period. This hypothesis is further supported by the evolutionary history and ecology of Isoetaceae which suggests they were prominent parts of the carboniferous flora with morphology comparable to extant *Isoëtes* species. Since NCA seems to be so evolutionarily plastic as to be better understood as a behavior, not a metabolic pathway, it will require a substantial future work to clarify the role of stomatal development, regulatory induction of NCA and atmospheric versus aquatic gas exchange in the genus Isoëtes, and other plants showing CAM-like biochemistry that are subject to evolutionary pressures other than drought stress.

## SUPPLEMENTARY DATA

### FUNDING

This work was supported by The Edmund Niles Huyck Preserve Graduate Research Grant in 2018. JSS is funded in part through the Department of Organismic and Evolutionary Biology graduate student research fellowships as well as the Arnold Arboretum fellowship program.

## ACKNOWLEDGEMENTS

The authors would like to thank Katharine E. Black, Sylvia P. Kinosian and Weston L. Testo for their helpful edits on the manuscript. We thank the Edmund Niles Huyck Preserve for funding, access to plant materials, and lodging during the field experiment, as well as the growth facilities staff at Weld Hill Research Building of the Arnold Arboretum of Harvard University for assisting in plant cultivation and growth chamber accommodations.

